# Water and the biology of prions and plaques

**DOI:** 10.1101/000224

**Authors:** Graham K. Steel, Philippa M. Wiggins

## Abstract

This is an attempt to account for the insolubility and/or aggregation of prions and plaques in terms of a model of water consisting of an equilibrium between high density and low density microdomains. Hydrophobic molecules, including proteins, accumulate selectively into stable populations, enriched in high density water, at charged sites on biopolymers. In enriched high density water, proteins are probably partially unfolded and may precipitate out when released. All extracellular matrices contain such charged polymers. Prions, which have been shown to accumulate in soils and clays containing silicates and aluminates also probably accumulate in extracellular matrices.

Release of proteins follows hydrolysis of the charged groups by highly reactive high density water. This is normally a slow process but is greatly accelerated by urea. Plaques may form with age and disease because of accumulation of urea and, perhaps, glucose in the blood. This favours precipitation of proteins emerging from matrices, rather than refolding and solution. Dialysis should, therefore, interfere with plaque formation and impede the development of some age-related diseases.

## Introduction

The search for the cause(s) of transmissible spongiform encephalopathies (TSEs) has a long and tortuous history. It is now widely accepted that nomenclature such as PrP^d^ and PrP^TSE^ denotes infectivity. Much of the “torture” surrounds whether Prions are the cause or the symptoms of TSE’s and also whether other particles are involved at some stage in the disease. In a recent paper^1^, 25-nm virus-like particles were found that were repeatedly found in cell cultures infected with Creutzfeldt-Jakob disease (CJD) and scrapie. Such particles are not only similar, but identical to virus-like tubulovesicular structures (TVS) found in experimental scrapie as early as in 1968, and subsequently not only in some, but in all naturally occurring and experimentally induced TSEs.

Whilst such particles have been generally been disregarded my most of the scientific community in this field mainly due to a poor track record of data although virus-like structures have been repeatedly reported got almost four decades in TSEs. TVS are spherical or tubular particles of approximate diameter 25–37 nm. They are smaller than synaptic vesicles, but larger than many particulate structures of the central nervous system, such as glycogen granules. Their electron density is higher compared with synaptic vesicles, and in experimental murine scrapie, they form paracrystalline arrays. Whilst what has been observed here over four decades cannot differentiate between TVS as a critical part of PrP^d^, or as a highly specific phenomenon, but their consistent presence in all not some TSEs clearly demands wider recognition.

## The role of water

Prions and plaques have hitherto been discussed in terms of the classical model of water, according to which molecules form an extensive randomly hydrogen-bonded network. Hydrogen bonds break and reform in picoseconds, their strength varying continuously from weak to strong. Observed properties of water (infrared spectra, free energies of hydration etc.) have been interpreted in terms of this model. Recently the model has been simplified. Now, according to Robinson and coworkers^2^ and Stanley and coworkers^3^ water exists as microdomains of high density water (HDW, density 1.2 g/ml) and microdomains of low density water (LDW, density 0.91 g/ml). As Nature frequently shows us, this apparent simplification has many and varied consequences, some highly complex. These microdomains must be in equilibrium, or the one of higher chemical potential would convert to the one with lower chemical potential. This equilibrium state of the two types of microdomains is responsible for most of the often unexpected consequences of this proposal, which has turned out to have great explanatory power in biology^4^^,^^5^. Some examples are: a mechanism for the prebiotic determination of the chiral specificity exhibited by all biopolymers; a mechanism for spontaneous synthesis of ATP (adenosine triphosphate), peptides etc.; a mechanism for mineralisation of bone; an explanation of the exquisite discrimination between sodium and potassium.

### Solutes and the mixture model

Solubility in the three-dimensional network of water molecules is simply a matter of balance between the forces operating to hold the ions or molecules of a solid together and the opposing forces of mixing those ions or molecules with water. The simplified model of just two types of hydrogen-bonding immediately reveals its extra dimensions. Since it is agreed that the properties of water are dominated by its mutual hydrogen-bonding of molecules, it must also be agreed that the two types of microdomain differ in all physical and chemical properties. The first property investigated was solvent power. It has turned out that all solutes investigated partition selectively either into LDW or into HDW. The immediate consequence of this is that any solute induces an osmotic pressure gradient between HDW and LDW, thus putting water out of equilibrium. For example glucose partitions selectively into LDW. The response of the system is to abolish the gradient by converting some LDW into HDW. Thus glucose is called a chaotrope because it induces HDW with increased entropy.

Glucose has only a modest preference for LDW, and the thermodynamic cost of displacing the HDW/LDW equilibrium, on top of the thermodynamic cost of separating glucose molecules from the solid, is easily met. If, however, a molecule has an extremely large preference for either type of microdomain, the thermodynamic cost of its solution is too great and that molecule is insoluble. For example ethane which partitions selectively into HDW has such a high preference that it is insoluble in the mixture. But ethanol, which replaces just one H with an OH, is highly soluble. So whether a molecule is soluble or not depends upon the mix of: moieties which partition into LDW and of moieties which partition into HDW.

This applies also to neutral salts. If both cation and anion induce LDW (eg CaSO_4_) solubility is low; if the cation induces LDW and the anion HDW (eg CaCl_2_) the net movement of the water equilibrium is slight and solubility extremely high.

### Multiple solutes

Since solubility depends upon the overall displacement of the water equilibrium, ethane could become soluble in the presence of another solute with a very great preference for LDW. In other words, solubility is not absolute and unchangeable it is a complex function of all components of a molecule and all components of a solution.

### Water and proteins

All proteins have both hydrophobic (LDW-inducing) and hydrophilic (HDW-inducing) moieties. Sometimes, however the excess of one over the other requires such a large displacement of the HDW/LDW equilibrium that the extended molecule is insoluble.

For example a highly hydrophobic protein in its extended state is insoluble, but it folds to remove much (not all) of that hydrophobic surface out of contact with water. Therefore the folded protein is soluble. Anything that prevents that folding will put the protein out of solution. Conversely if the other components of the solution are changed, the unfolded protein may be able to dissolve. For example urea at high enough concentrations unfolds (denatures) hydrophobic proteins and allows them to remain in solution in their extended state. Similarly a protein with a large excess of hydrophilic amino acids may be insoluble in its extended state but can fold to reduce the area of contact between water and some hydrophilic groups, and become soluble. These are mechanisms which can force a protein out of solution, or solubilise an insoluble protein. Most biological small molecules are mixtures of HDW-inducing and LDW-inducing moieties. They are, therefore extremely soluble.

### Solutes which might be expected to solubilise hydrophobic molecules

In order to decrease the extent to which LDW-inducing molecules such as ethane displace the HDW/LDW equilibrium, all that is needed is a cosolute which has an extreme preference for LDW and consequent tendency to induce HDW. These solutes have been identified: they include: H_2_PO_4_^-^, HSO_4_^-^, NO_3_^-^, HCO_3_^-^, I^-^, Br^-^, Cl^-^ and NH_4_^+^, Cs^+^, Rb^+^, K^+^. Non-electrolytes which are much less effective include urea and glucose. The effectiveness of ions appears to follow their size, the larger ions being more effective.

### Compatible surfaces

A folded protein, then, is soluble because it has reduced the area of contact between its hydrophobic components and water. Most surfaces, can only modify their area of contact with water by adsorbing molecules from solution. Therefore surfaces participate in the many factors which determine the position of the HDW/LDW equilibrium in solution. This becomes of particular importance in biology where vessels are narrow and cells extremely small and crowded and no solute is ever far from a surface. *In vivo* surfaces, being made of proteins and polysaccharides, are, like glucose, neither excessively HDW-inducing nor excessively LDW-inducing. Therefore they have little tendency to adsorb molecules or cells from solution. For example red cells squeeze through tiny arterioles without sticking to sides. These are called compatible surfaces. A foreign surface, however, such as glass (hydrophilic) or plastic (hydrophobic) does have an effect on the water equilibrium and attracts molecules and cells out of solution or suspension to lower its surface area of contact with water. This was demonstrated with a series of experiments using hydrophilic and hydrophobic (methylated) glass beads and hydrophilic and hydrophobic surfaces. Hydrophilic beads stuck to both hydrophilic and hydrophobic surfaces, and hydrophobic beads also stuck to both hydrophilic and hydrophobic surfaces^5^. The stickiness of these surfaces could obviously not be attributed to binding sites. It illustrated that ‘binding’ could sometimes be driven by the subtle thermodynamic costs of displacing the HDW/LDW equilibrium, without specific bonds of any kind. In similar experiments with cells in suspension, all cells (red cells, platelets, stem cells) stuck to glass and stuck to plastic and lysed, but survived when contained in compatible surfaces.

### Plaque formation

Once, for whatever reason, a few proteins have precipitated, they present to the solution a hydrophobic incompatible patch of surface on to which other polymers and cells will non-specifically stick to reduce the surface/water interface; the proteins will unfold leading to plaque, fibril and tangle formation.

### The solubility of prions

Prions are evidently only very slightly soluble and quite readily become insoluble. They must have a slight solubility or they would not be able to enter the bloodstream. Movement of molecules from one site in the body to another absolutely requires solution in water or association with other soluble molecules. Their slight solubility in water, which is attributed to their hydrophobicity, is, in fact, a consequence, not of hatred of water, but of an inordinate affinity for HDW. The universally used term hydrophobicity is a misnomer.

### Prions and charged polymers such as heparan and dextran sulphate

*In vivo* prions can exploit their high affinity for HDW because stable populations of enriched HDW and enriched LDW coexist in gels and matrices with charged groups such as -COO^-^, -HSO_3_^-^, -HPO_4_^-^, -NH_3_^+^. These populations have high capacities for accumulating solutes from the bulk solution. The mechanism of generation of HDW in such matrices is illustrated in Figure 1. The populations are represented as pure HDW and pure LDW. In fact they are enriched in one type or the other rather than absolutely pure HDW or LDW. The negative charge on the surface of a polymer (eg heparan sulphate or dextran sulphate) is balanced by a cation in solution. Across the imaginary interface between the zone containing the cation and the rest of the solution, there is a substantial osmotic pressure gradient. (a single ion in a few nm^3^ of water translates into a molar concentration). Water cannot move in to abolish this gradient because of the powerful attraction between positive and negative charges. The tendency of water so to move, however, becomes a positive pressure acting on water inside that zone and a negative pressure on water just outside the zone. This converts some water immediately adjacent to the surface to HDW and some of a zone of water outside that to LDW. Similarly a positively charged polymer induces stable populations of HDW at the surface and LDW further out. The sizes of these zones are determined by the attraction between fixed charge and counterion, the other components of the solution and the specific counter ion. They are, in general 1-4 nm across. In Figure 1 a folded prion is selectively taken up into the HDW.

**Figure 1.**
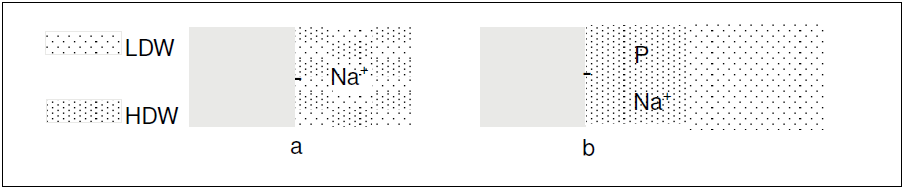
a, a negatively charged group on a polymer is neutralised by a Na^+^ ion in solution in the mixture of waters, generating an osmotic pressure gradient between that zone of water and the rest of the solution; b, the powerful force between the two charges prevents entry of water to abolish the gradient but it acts a pressure gradient across the imaginary interface between water at the surface and the rest of the water. Positive pressure on water at the surface converts it to HDW while negative pressure converts an outside zone into LDW. The hydrophobic prion (P) selectively partitions into HDW.

Van Horssen et al^7^ has shown that heparan sulphate is associated with Alzheimer plaques. It is suggested that part of their role involves selective accumulation of hydrophobic proteins into HDW. In general extracellular matrices consist of such polymers carrying many both positive and negative charges. Moreover, prions are found in soils and clays which also carry charged silicates and aluminates^8^. Therefore prions from their low concentration in blood may be accumulated to high concentrations in pockets of HDW. Extracellular matrices comprising mostly charged polysaccharides occur in most tissues and have been shown to be storage sites for hormones, growth factors etc. If they selectively accumulate prions into their HDW, they can also be reservoirs for prions. This makes a significant difference to the biology of prions, which, mainly held in the numerous extracellular matrices, could still exist as extremely low concentrations in the blood. These low blood concentrations would be in equilibrium with the large majority of prions in the matrices. An important property of the matrix prions is that they exist in a more extended state than in the water mixture. This follows because the driving force for folding of prions in the mixture of waters was the thermodynamic cost of displacing the HDW/LDW equilibrium, as the extended molecule induced so much LDW. With the molecule in solution in enriched HDW, this driving force for folding is much weaker.

### Leakage of prions from extracellular matrices

HDW is extremely reactive^4^. Water as a mixture of microdomains is prevented from ionising freely because the production of H^+^ + OH^-^ (both powerful inducers of LDW) requires too great a displacement of the water equilibrium.

An enriched pocket of HDW, however, has less constraint. It behaves as a strong acid and strong base together. Therefore, relatively slowly, because a covalent bond has to be broken, the stable pocket of HDW illustrated in Figure 1 collapses as the fixed negative charge is hydrolysed off the surface (Figure 2). The prion emerges into the blood in its relatively extended form which is insoluble. It then, either precipitates out before there is time to fold, aggregates with its fellows, or folds and goes into solution and is available to be taken up into other pockets of HDW. This steady leakage of prions, soluble and insoluble, gives the appearance of proliferation.

**Figure 2.**
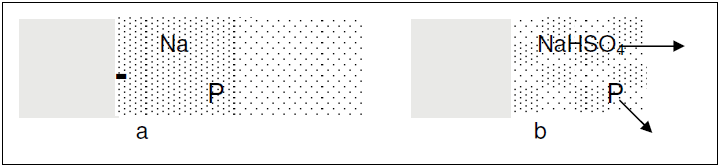
a, negative charge fixed to the surface and neutralised by a sodium ion; b, HDW has hydrolysed the negative charge, water reverts to the mixture of HDW and LDW and P and NaHSO_4_ diffuse out.

### A difference between prions and plaques

Prions are invariably fatal at any age. Plaques, on the other hand, appear only with age or with disease. i.e. something happens with age or disease that makes a normally soluble protein precipitate out. In youth and health normally functioning kidneys maintain a constant composition of the blood and interstitial solution. Plaques, then, probably appear only when impaired kidney function relaxes its tight control of solutes. As previously mentioned, some consequences of the mixture model of water are unexpected and complex. One of these is the effects of small solutes on the isolated populations of HDW and LDW illustrated in Figures 1 and 2. See Figure 3.

**Figure 3.**
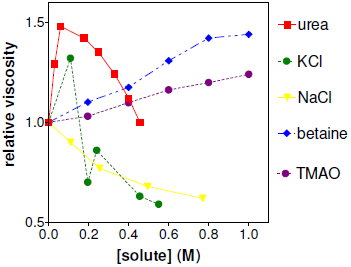
Effects of added solutes on the viscosity of a dextran sulphate solution (1 g dextransulphate in 3 g water.)

Urea and KCl, in particular, illustrate these complexities. These results are discussed in more detail elsewhere^4^, but are of present interest because urea is one of the likely candidates for increased blood concentration with age and disease; it is generally a chaotrope, inducing HDW. In Figure 3, however, it increases viscosity (i.e. increases LDW) up to 0.05 M and then decreases viscosity above 0.4 M (not shown in this graph). Linearity with concentration is one of the many things that must be abandoned in this version of water. Moreover, in spite of the apparent simplicity of a regime with just two strengths of hydrogen bonds, it is usually impossible to ratonalise a single experiment. More information is needed. Urea in Figure 3 is an example. In other unpublished experiments, urea, at quite low concentrations, greatly accelerated the release of silicates from glass beads, the release of NH_4_^+^ from an anion-exchange resin and release of HSO_4_^-^ from a cation-exchange resin. In other words urea has effects which could not be deduced only from its effects upon the viscosity of dextansulphate solution, unexpected as those effects are Combining these different consequences of urea, however, allows a plausible explanation of its effects. This may be relevant to the occurrence of plaques only with age or disease. Apparently (see Figure 4) urea goes selectively into LDW and water moves from the HDW zone to abolish the osmotic pressure gradient. This has the effect of increasing the amount of LDW and therefore viscosity of dextran sulphate.

The only way to explain the increase in hydrolysis of the charged group is that the volume of HDW surrounding the fixed charge has decreased so that the excess concentration of the counter ion has increased and the pressure gradient must also increase. Therefore the water adjacent the charged group is further enriched in HDW and is accordingly more active. Any solute that decreases the volume of HDW will have the same effect. Butanol and ethanol both accelerate hydrolysis. They accumulate into HDW and, since they decrease the dielectric constant, they allow the counter ion to move closer to the fixed charge. This suggests that prions and other hydrophobic proteins will also accelerate hydrolysis, but urea must accelerate it yet more.

**Figure 4.**
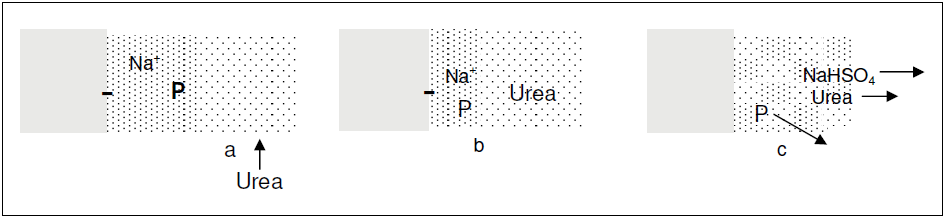
a, Low concentrations of urea accumulate into LDW; b, water moves from HDW to LDW to abolish the osmotic pressure gradient. This increases LDW and decreases HDW. But, as the distance between fixed charge and counterion decreases, the excess concentration of the counter ion increases. This increases the pressure gradient and further enriches the zone in HDW; c, The fixed charge is prematurely hydrolysed. And all solutes diffuse out.

### Plaques and prevention of plaques

A possibility to be considered, therefore, is that extracellular matrices are always reservoirs for hydrophobic proteins. While there, the proteins do no damage, and as they are slowly released they can complete folding and dissolve, to be taken up by other matrices. If, however, the blood contains a higher-than-normal concentration of urea or glucose (in type 2 diabetes) the rate of release becomes too great for refolding before solution, and the protein precipitates out. The consequence is a series of patches of incompatible surface to which other proteins, polysaccharides, cells, non-specifically stick, giving plaques and tangles and ensuring continuing precipitation.

### Effects of precipitated prions and plaques

One effect of this precipitation may resemble the role of cholesterol in coronary arteries. The presence of an insoluble substance sticking to the surface, narrows arteries and arterioles and impairs blood flow to organs. It seems unlikely that the soluble form of the prion or plaque is the cause of malfunction, so that therapeutic measures should concentrate on either preventing accumulation of hydrophobic proteins into matrices, or better, controlling their release. It is probably not possible to prevent their accumulation, but their rate of release may well be amenable to control. There must be many plaques which are not damaging, just as there may be cholesterol deposits which are not damaging. Only those deposits which significantly narrow or block important vessels are likely to contribute to the observed pathology. There may be time, therefore, for rescue, if at the earliest symptoms the patient is dialysed to remove excess urea and/or glucose and prevent further rapid release and precipitation of proteins. The alternative, of course is to oppose the effect of urea. NaCl is already doing this. NaCl increases the local dielectric constant and allows the counter ion to move away from the fixed charge. This decreases the osmotic gradient and delays hydrolysis of the fixed charge group. Lithium would be the best candidate to oppose urea. It is much safer, however, to remove harmful solutes than add others, so probably early dialysis is the better treatment for plaque formation. Prions may respond to lithium, which, added in low concentrations should accumulate into HDW and supplement the role of NaCl.

Prions, however, have been shown to cross cell membranes^9^^,^ ^10^. The potential damage of intracellular precipitation of proteins is extremely high. Apart from low concentrations of ions, metabolites and water, cells are polyelectolytes (proteins, DNA, RNA, polysaccharides). The damaging sequence of accumulation in HDW at positive and negative charged sites, minimal folding, cleavage of the charged group from the matrix and release of the minimally-folded prion must continue unabated in the fragile intracellular environment, with lethal consequences.

